# Splice Expression Variation Analysis (SEVA) for Inter-tumor Heterogeneity of Gene Isoform Usage in Cancer

**DOI:** 10.1101/091637

**Authors:** Bahman Afsari, Theresa Guo, Michael Considine, Liliana Florea, Luciane T. Kagohara, Genevieve L. Stein-O’Brien, Dylan Kelley, Emily Flam, Kristina D. Zambo, Patrick K. Ha, Donald Geman, Michael F. Ochs, Joseph A. Califano, Daria A. Gaykalova, Alexander V. Favorov, Elana J. Fertig

**Affiliations:** Department of Oncology, Division of Biostatistics and Bioinformatics, Sidney Kimmel Comprehensive Cancer Center, Johns Hopkins University, Baltimore, MD, 21205, USA; Department of Otolaryngology - Head and Neck Surgery, Johns Hopkins University, Baltimore, MD, 21205, USA; McKusick-Nathans Institute of Genetic Medicine, Johns Hopkins University, Baltimore, MD, 21205, USA; Department of Otolaryngology – Head and Neck Surgery, University of California, San Francisco, CA, 94158, USA; Department of Applied Mathematics & Statistics, Johns Hopkins University, Baltimore, MD, 21218, USA; Department of Mathematics & Statistics, The College of New Jersey, Ewing, NJ, 08628, USA; Division of Otolaryngology, Department of Surgery, University of California, San Diego, CA, 92093, USA; Laboratory of Systems Biology and Computational Genetics, Vavilov Institute of General Genetics, RAS, Moscow, 119333, Russia; Laboratory of Bioinformatics, Research Institute of Genetics and Selection of Industrial Microorganisms, Moscow, 117545, Russia

## Abstract

**Motivation:** Current bioinformatics methods to detect changes in gene isoform usage in distinct phenotypes compare the relative expected isoform usage in phenotypes. These statistics model differences in isoform usage in normal tissues, which have stable regulation of gene splicing. Pathological conditions, such as cancer, can have broken regulation of splicing that increases the heterogeneity of the expression of splice variants. Inferring events with such differential heterogeneity in gene isoform usage requires new statistical approaches.

**Results:** We introduce Splice Expression Variability Analysis (SEVA) to model increased heterogeneity of splice variant usage between conditions (e.g., tumor and normal samples). SEVA uses a rank-based multivariate statistic that compares the variability of junction expression profiles within one condition to the variability within another. Simulated data show that SEVA is unique in modeling heterogeneity of gene isoform usage, and benchmark SEVA’s performance against EBSeq, DiffSplice, and rMATS that model differential isoform usage instead of heterogeneity. We confirm the accuracy of SEVAin identifying known splice variants in head and neck cancer and perform cross-study validation of novel splice variants. A novel comparison of splice variant heterogeneity between subtypes of head and neck cancer demonstrated unanticipated similarity between the heterogeneity of gene isoform usage in HPV-positive and HPV-negative subtypes and anticipated increased heterogeneity among HPV-negative samples with mutations in genes that regulate the splice variant machinery.

**Conclusion:** These results show that SEVA accurately models differential heterogeneity of gene isoform usage from RNA-seq data.

**Availability:** SEVA is implemented in the R/Bioconductor package GSReg.

## 1 Introduction

Alternative splicing events (ASE) are biological mechanisms that enable expression of a variable repertoire of protein products from a single protein coding gene (Lim *et al.*, 2011). Differences between the resulting protein products associated with cell types identity, cellular functions, and phenotypes. In alternative splicing, distinct sets of RNA fragments (exons and even introns) within a single gene are included into mature mRNA. Each unique set of exons and introns is called a gene isoform, and need not contain canonical adjacent exons within a mRNA molecule (reviewed in Keren *et al.* 2010). Cells contain a splicing machinery complex to regulate which gene isoforms are produced, and cause the expression of specific gene isoforms to be under tight control within healthy cells, accounting for both their identity and environmental context. Numerous bioinformatics algorithms have been developed for analysis of alternative splice variation from RNA-seq data. Reads in RNA-seq data may span multiple exons and introns, thereby providing evidence for expression of specific gene isoforms. Bioinformatics analysis of RNA-seq data can identify the expression of gene isoforms in a single sample (Song *et al.*, 2016; Canzar *et al.*, 2016; Pertea *et al.*, 2015; Guttman *et al.*, 2010; Li *et al.*, 2011;Wang *et al.*, 2010). The resulting isoform expression profiles from different samples can be further aggregated to quantify gene isoform usage. These data are input to methods for differential ASE analysis that compare relative changes of expected expression of isoforms between phenotypes to define the differences in the landscape of gene isoform usage between phenotypes (Li *et al.*, 2011; Anders *et al.*, 2012; Hu *et al.*, 2013; Shen *et al.*, 2012, 2014; Leng *et al.*, 2013). These methods model isoform usage that is consistent between samples from the same phenotype and differs consistently with isoform usage in samples from another phenotype. Thus, their underlying algorithms assume that the splicing machinery works properly in both phenotypes. They also assume that mixture of cell types and, thus, the repertoire of gene isoforms, is similar within samples from the same phenotype.

Changes to splicing events are pervasive in cancer and can be used as biomarkers (Sebestyen *et al.*, 2015). In contrast to normal samples, the splice variant machinery is frequently altered or broken in tumors (Ebert and Bernard, 2011). In addition, tumors are comprised of a heterogeneous mixture of cell types, each of which has a distinct milieu of expressed gene isoforms. Both of these factors result in more variable gene isoform usage within tumor samples relative than in normal samples (Li *et al.*, 2014; Guo *et al.*, 2017). Notably, differential ASE methods are not designed to model these differences in heterogeneity. Instead, they model differences in the expected relative expression levels instead of their relative heterogeneity. Detection of alternative splicing in tumors modeling differential heterogeneity of gene isoform usage in tumors requires a new bioinformatics algorithm.

In this paper, we develop a novel algorithm called Splice Expression Variability Analysis (SEVA) for differential heterogeneity of gene isoform usage between samples from two phenotypes. This algorithm uses a multivariate, non-parametric statistic of the heterogeneity of expression profiles for gene isoforms in tumor relative to normal samples. The measure of heterogeneity is adapted from the Kendall-tau dissimilarity measure and is applied to compare the junction expression profiles between all pairs of samples of each phenotype for each gene. Junction expression quantifies the number of reads that span a pair of exons or an exon to an intron, thereby providing direct evidence for the gene isoform expression (Fig 1). Thus, SEVA can provide both a measure of heterogeneity of alternative splicing events in each phenotype and a statistic for the significance of the difference of these measures in two phenotypes. SEVA is implemented as a new function in the R/Bioconductor package GSReg (Afsari *et al.*, 2014b). SEVA is a natural extension of this package, which is devoted to differential variation of gene regulation in cancer and previously implemented algorithms for pathway dysregulation (Eddy *et al.*, 2010; Afsari *et al.*, 2014b).

**Figure 1:**
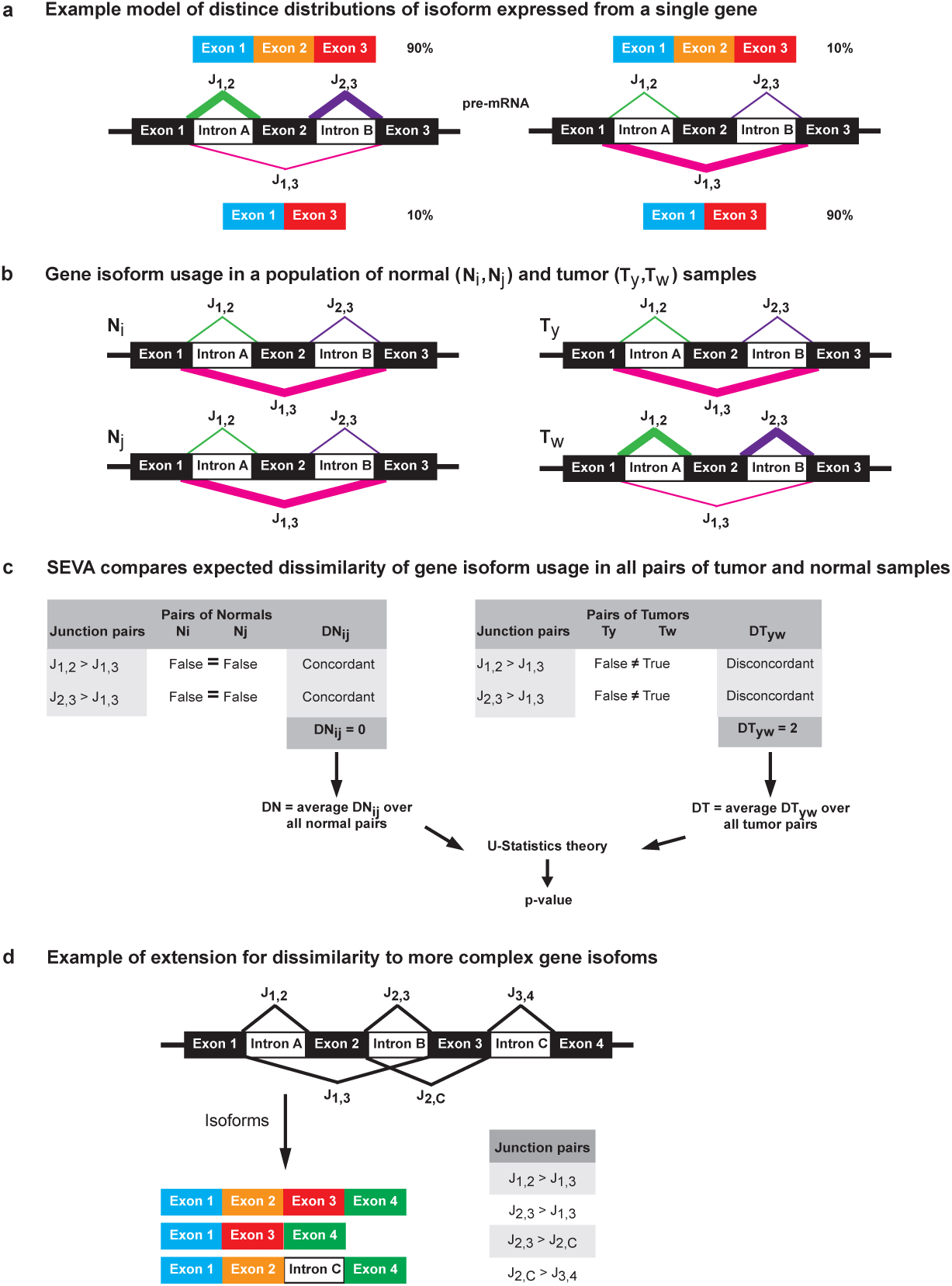
Overview of SEVA. (a) Relative junction expression quantifies the distribution of isoform usage of a gene. For simplicity of this example, we show a gene with three exons. The model is shown for two samples of gene isoform usage: one with higher relative expression of an isoform with all three exons (left) and another with higher relative expression of an isoform that skips the middle exon (right). The relative strength of junction expression in overlapping pairs (e.g., *J*_1,2_ with *J*_1,3_ or *J*_2,3_ with *J*_1,3_) corresponds to the relative proportion of isoform usage. (b) Example of gene model from (a) in multiple normal (N, left) and tumor (T, right) samples. Note that the normal samples have lower heterogeneity of gene isoform usage than the tumor samples. (c) To quantify isoform expression, SEVA compares the expression of all pairs of overlapping junctions (see a and d). A dissimilarity measure is obtained from the concordance of the comparisons of pairs of overlapping junctions in each pair of samples. This measure is applied to all pairs of samples from the same phenotype (see b) and then U-statistics theory is applied to these measures to compare the variation of gene isoform usage between the phenotypes. (d) Extension of (a) for a more complex gene splicing model.

## 2 Methods

### 2.1 HNSCC RNA-seq datasets

We use RNA-seq data for 46 HPV+ HNSCC and 25 independent normal samples from uvulopalatopharyngoplasty previously described in Guo *et al.*, 2016 and 44 HPV+ HNSCC, 235 HPV-HNSCC, and 16 matched normal tissues from the freeze set for TCGA HNSCC (Cancer Genome Atlas Network, 2015).

### 2.2 *In silico* data

We base our simulated data on the isoform expression from RNA-seq data for normal samples from Guo *et al.* 2016. To reduce the size of the simulations, we limit the analysis to 600 preselected genes in chromosome 1. We require these genes to have mean log2 expression gene counts from the TCGA RSEM v2 pipeline Cancer Genome Atlas Network (2015) between 4 and 9 and to have at least 2 isoforms. This filtering criterion ensures that genes have differential gene isoform usage because they are both expressed and have alternative splice variants. We generate a simulated dataset of 25 tumor and 25 normal samples as follows. To simulate normal samples, we calculate the average isoform expression for these genes in real normal samples and input these values to Polyester with default parameters (Frazee *et al.*, 2015b) to generate simulated RNA-seq reads.

We then simulate alternative splicing and differential expression events in tumor samples for a pre-selected subset of the 600 genes, with 150 differentially spliced, 150 differentially expressed, 150 both, and 150 neither. The expected isoform expression of a gene with alternative splicing in a tumor sample is obtained by randomly permuting the expected expression of all isoforms for that gene in the normal samples (Supplemental Fig 1). Such random distribution of expected isoform expression is similar to previous simulation studies which randomly distributing expression values among the set of all isoforms of a gene (Alamancos *et al.*, 2015; Liu *et al.*, 2014). A gene is simulated to have differential expression by either doubling the expression of all its isoforms relative to the values in normal samples or halving the expression of all its isoforms, where doubling (over-expressed) or halving(under-expressed) was selected at random.

The simulation models tumor heterogeneity of gene isoform usage by varying the number of tumor samples with alternative splice variant usage and differential expression. One subset of tumor samples, called normal-like tumors, follows the same distribution of isoform expression as normal samples. The other subset follows the distribution of isoform expression for tumor samples (Supplemental Fig 2). The resulting values for expected isoform distribution in each subtype are input to Polyester (Frazee *et al.*, 2015b) to generate simulated RNA-seq reads for tumor samples. In total, four simulated datasets are created by varying this parameter. Each set had 25 simulated normal and 25 simulated tumor samples. We create an MDS plot of one of the differentially spliced genes (identified by SEVA in a cohort with 10 normal-like tumor samples) and note the similarity of this MDS plot to similar plots in real data described in the results (Section 4.3).

Simulated reads for both normal and tumor samples are normalized as described in the data normalization subsection below.

### 2.3 RNA-sequencing data normalization and mutation calls

All *in silico* and real RNA-seq data are normalized with the RNA-seq v2 pipeline from TCGA (Cancer Genome Atlas Network, 2015). Junction expression is obtained directly from the Map-Splice (UNIX multithread version 2.0.1.9) (Wang *et al.*, 2010) output for each sample, setting expression values to zero for junctions that are not detected from MapSplice in a given sample. Simulated data is also aligned with TopHat2 version 2.1.0 (Kim *et al.*, 2013). Gene and isoform expression data from TCGA are obtained from the level 3 normalized data, but junction expression is obtained by rerunning MapSplice to perform *de novo* junction identification to compare with the training RNA-seq data from (Guo *et al.*, 2016). Comparisons between HPV-positive and HPV-negative HNSCC are made for level 3 junction data for previously annotated junctions for TCGA samples available on FireBrowse. TCGA samples with mutations or copy number alterations any of the *SF3B1, SF1, SF3A1, SRSF2, U2AF1, U2AF2, ZRSR2*, or *PRPF40B* genes in cBioPortal (Gao *et al.*, 2013) are said to have altered RNA splice machinery based upon annotations in Ebert and Bernard, 2011.

### 2.4 Implementation and software

SEVA is implemented in the R/Bioconductor package GSReg (version 1.9.2) (Afsari *et al.*, 2014a). Junctions are assigned to a gene using the UCSC gene annotations in the R/Bioconductor package TxDb.Hsapiens.UCSC.hg19.knownGene.db (version 3.2.2). The analyses presented in this study removes genes with only one junction from analysis. Additional filtering criteria are described in the vignette, but not used. The SEVA analysis of junction expression is computationally efficient. All of the SEVA computations for simulated data completed in less than an hour for simulated data, less than an hour for the46 tumors and 25 normals in the HPV+ HNSCC cohort from Guo *et al.* (2016), and less than two hours for the 279 HNSCC tumors in TCGA (Cancer Genome Atlas Network, 2015) on a MacBook Pro with Core (TM) i7-3720QM Intel CPU @2.6 GHz. Genes with Benjamini-Hochberg adjusted p-values below 1% are statistically significant. All code for the SEVA analyses is available from https://github.com/FertigLab/SEVA. In the Supplemental Methods, we show that the computations for SEVA grows as the cube of the sample numbers and linearly with the number of genes, with a dependence upon the number of overlapping junctions per gene.

EBSeq is performed with the R/Bioconductor package EBSeq version 3.3 (Leng *et al.*, 2013). Isoform expression for all genes is the input in the EBSeq analysis. Isoforms with posterior probability above 99% are called significantly differentially spliced. EBSeq is also applied to gene expression values, and genes with a posterior probability above 99% are significantly differentially expressed. DiffSplice 0.1.2 beta version (Hu *et al.*, 2013) is run directly on aligned RNA-seq data obtained from the MapSplice alignment. Default parameters are used, with a false discovery rate of 0.01. Because DiffSplice requires equal numbers of samples in each group, we select a random subset of 14 HPV+ HNSCC and 14 normal samples from the dataset in (Guo *et al.*, 2016). rMATS version 3.2.5 (Shen *et al.*, 2014) is run on TopHat2 (Kim *et al.*, 2013) aligned simulated data. To produce read counts to use as reference annotation for rMATS, we constructed a merged gene annotation set by running cuffmerge v2.2.1 on the transcript predictions from the individual samples, produced with CLASS2 v.2.1.6 (Song *et al.*, 2016),and the hg19 gene annotations used in the TCGA RSEM v2 pipeline (Cancer Genome Atlas Network, 2015). Read counts are generated with Ballgown tablemaker v.2.1.1 (Frazee *et al.*, 2015a).

We perform cross study validation by comparing whether statistics in one cohort are significantly enriched in the other using the function wilcoxGST in LIMMA version 3.24.15 (Ritchie *et al.*, 2015).

## 3 Algorithm

### 3.1 Splice Expression Variation Analysis (SEVA)

A gene can have multiple isoforms. RNA-seq data provide direct evidence for the expression of a specific gene isoform based upon coverage of the junctions in that isoform (Fig 1a,d). Such junction expression can also indicate the simultaneous expression of multiple isoforms in a sample (Fig 1a,b). If the distribution of isoform expression changes between samples, then so too does the distribution of the expression for the set of all junctions in the gene. Thus, the multivariate distribution (i.e., the joint distribution) of the set of junction expressions can quantify gene-level changes in isoform usage between sample groups. In this study, we hypothesize that the expression of ASE variants is more heterogeneous than the expression in normal samples (Fig 1c). We develop a new method called Splice Expression Variability Analysis (SEVA) that compares the variability of the multivariate distribution of junction expression profiles between phenotypes to test the hypothesis of differential heterogeneity of gene isoform usage.

The key observation that underlies SEVA is that junctions which correspond to distinct isoforms are overlapping, and therefore mutually exclusive. Changes in the relative expression of gene isoforms between a pair of samples result in a corresponding switch in the relative expression of specific junctions. Namely, the switching occurs in the overlapping junctions. An example of this switch is shown for a simple exon skipping event in Fig 1a. In this case, the expression of the junction between the first and third exon corresponds to the switches with the expression of both the junctions in the isoform without the skipping event (between the first and second exon and between the second and third exon). This concept extends to more complex gene models (Fig 1d).

Based on the observation described above, SEVA defines a “splice dissimilarity” measure by computing the frequency of switching between all pairs of mutually exclusive junctions (Fig 1c). A switch occurs when the rank comparison between two overlapping junctions differs between samples. We note that Kendall-tau dissimilarity measure quantifying such discordance between the ranks of pairs of observations. Whereas Kendall-tau quantifies the rank discordances between all pairs of observations, SEVA limits this comparison to the mutually exclusive junction coverage. Such comparisons are made between all pairs of samples in the same phenotype to estimate the heterogeneity of gene isoform usage in that phenotype. Then, the algorithm tests for a significant difference in heterogeneity between the two phenotypes using an approximation from U-Statistics theory (Vaart, 1998)(Fig 1c, Supplemental Methods).

## 4 Results

### 4.1 SEVA accurately detects differential heterogeneity in ASE usage

We generate *in silico* RNA-seq data to benchmark the performance of SEVA relative to EBSeq (Leng *et al.*, 2013), DiffSplice (Hu *et al.*, 2013), and rMATS (Shen *et al.*, 2014) in detecting genes with known differential heterogeneity of isoform usage in populations of simulated tumor and normal samples (described in methods). The simulated dataset contains 25 normal and 25 tumor samples. The simulation generates four cohorts of tumor samples, which have 10, 15, 20, or 25 samples with differential gene isoform usage events. The remaining samples follow the distribution of gene isoform usage in normal samples (see Methods, Supplemental Figs 1 and 2). The simulation with 25 samples with differential gene isoform usage in tumor samples has the greatest mean difference relative to normal samples and the least heterogeneity of gene isoform usage within the tumor cohort. The simulations with 10 and 15 samples with differential gene isoform usage have the greatest heterogeneity. We compare the results of all algorithms to the true events in the four simulations to estimate precision (positive predictive value) and recall (sensitivity) (Fig 2).

**Figure 2:**
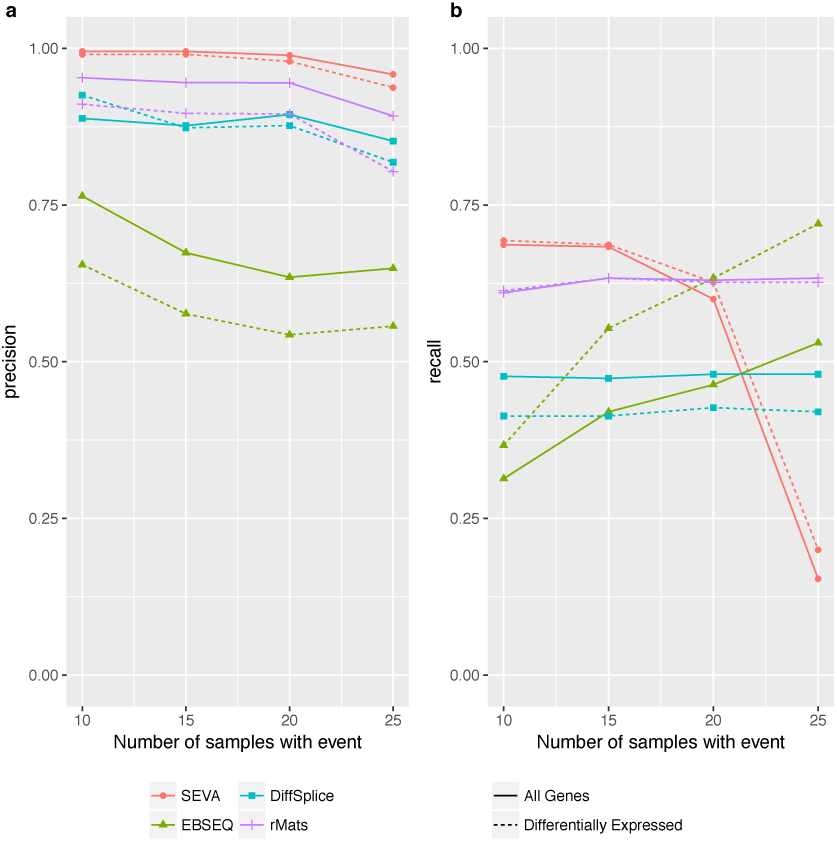
Performance in simulated RNA-Seq data. (a) Precision of different algorithms (in legend) on the simulated dataset. Varying numbers of the total tumor samples with alteration events (x-axis), with tumor heterogeneity decreasing along the x-axis. (b) Recall of simulated data, as in (a). Precision and recall computed separately for all genes(solid) and for the subset of 300 genes that are differentially expressed (dashed).

SEVA’s precision remains around 95% while that of DiffSplice fluctuates around 90% and the range of EBSeq’s precision is 60%-80%. The precision for rMATS is around 90% for differentially spliced genes that are not differentially expressed and ranges from 75% to 90% for differentially spliced genes that are also differentially expressed. These results are independent of the number of cancer samples containing the alternative gene isoform expression (Fig 2a). The precision of both SEVAand DiffSplice is independent of whether the gene is differentially expressed in addition to differentially spliced. EBSeq has lower precision for detecting differential splice status among differentially expressed genes compared to the precision in genes that are both differentially expressed and differentially spliced.

SEVA has the highest recall in the simulated dataset with greatest heterogeneity in gene isoform usage (fewer than 20 of the tumor samples), but drops sharply in the more homogeneous population of 25 tumor samples all containing the same gene isoform usage (Fig 2b). SEVA’s inability to detect alternative splicing events in homogeneous populations is consistent with the design of the algorithm. The recall for EBSeq remains consistently higher among genes that are both differentially expressed and differentially spliced than among genes that are only differentially spliced. Its recall increases with the number of tumor samples containing the alternative isoform usage for both types of genes. The recall for rMATS remains independent of the sample size and differential expression status of the genes. Both DiffSplice and rMATS have modest recall independent of tumor heterogeneity in the simulations. The precision of SEVA is independent of the number of isoforms in each gene, whereas recall is lowest for genes with only two isoforms and insensitive thereafter (Supplemental Fig 3). These results confirm that SEVA specifically identifies genes with high relative heterogeneity of gene isoform usage between sample phenotypes, whereas other algorithms identify genes with homogeneous differential gene isoform usage.

### 4.2 SEVA identifies a set of ASEs with variable gene isoform usage in 46 HPV+ HNSCC tumor samples relative to 25 normal samples

We use RNA-seq data for 46 HPV+ HNSCC and 25 normal samples from Guo *et al.*, 2016 as a benchmark for empirical analysis of SEVA in real sequencing data. SEVA identified 985 genes as having significant alternative gene isoform usage in cancer (compared to 2439 in EBSeq and 2535 in DiffSplice). In addition to identifying the altered gene isoforms in each class (e.g., tumors or normals), the statistics underlying SEVA enable quantification of relative variation in isoform usage for each gene in each of the phenotypes that are compared in the analysis. We plot these statistics to compare variation of isoform usage in all genes and the genes that SEVA calls statistically significant to test our central hypothesis that gene isoform usage is more variable in tumor than normal samples (Fig 3a). Since the ground truth is unknown in these real data, we cannot benchmark SEVA’s performance relative to other algorithms in terms of precision and recall. Nonetheless, consistent with our hypothesis, the variation in all genes is shifted towards higher variation in tumor samples. Moreover, more of these significant genes have more variable gene isoform usage in tumor samples than in normal samples (Fig 3).

**Figure 3:**
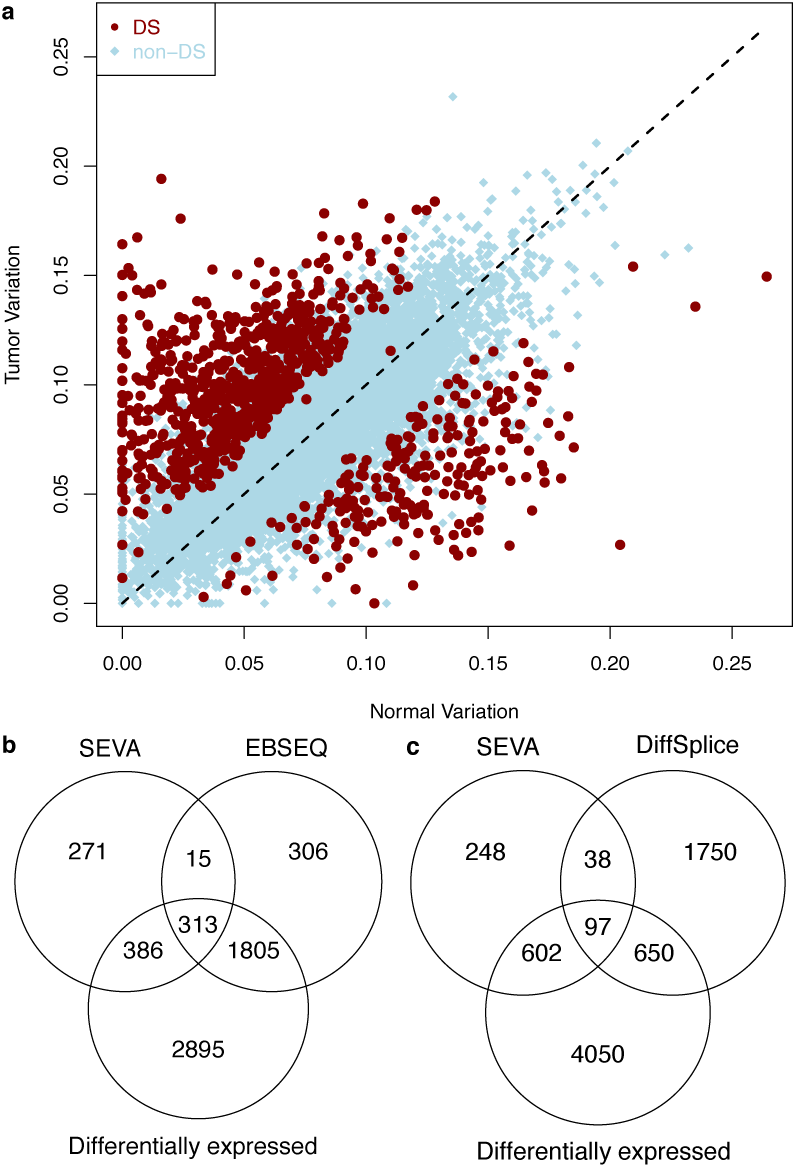
Comparison of splice variant events identified in different algorithms in real HPV+ HNSCC RNA-seq data. Variability of junction expression profiles corresponding to gene isoforms. Each point representsa gene, x-axis and y-axis its variability computed for SEVA in normal vs cancer, respectively. The red points represent significantly differentially spliced (DS) genes identified with SEVA, and blue genes that were not significantly spliced (non-DS). (b) Venn diagram comparing differentially spliced genes identified by SEVA and EBSeq, as well as differential expression status of each gene. (c) Comparison of SEVA and DiffSplice as described in (b).

### 4.3 SEVA analysis finds greater variation in tumor than normal samples in previously validated HNSCC-specific splice variants *TP63, LAMA3*, and *DST*

Recent data suggest that the majority of ASE in HNSCC (39%) are classified as alternative start sites (Guo *et al.*, 2017), which can be recognized by ASE-detection algorithms as insertion and/or deletion alternative splicing events. Indeed, alternative start site splice events in six genes (*VEGFC, DST, LAMA3, SDHA, TP63*, and *RASIP1*) were recently observed as being unique to HNSCC samples from microarray data (Li *et al.*, 2014). Three of these genes (*DST, LAMA3*, and *TP63*) were also confirmed as differentially spliced in HNSCC tumors with experimental validation in an independent cohort of samples, while the other three genes (*VEGFC, SDHA*, and *RASIP1*) were not confirmed (Li *et al.*, 2014). SEVA identified all three validated genes: *DST* (p-value 3,×10^−10^), *LAMA3* (p-value 1×10^−10^), and *TP63* (p-value 6×10^−10^) genes, as well as *RASIP1* (5×10^−7^) as genes having significant differential heterogeneity of isoform usage. Splicing of *SDHA* and *VEGFC* were found non-significant in agreement with experiments. Notably, EBSeq only identifies VEGFC to be differentially spliced (p-value 3 × 10^−6^) that was not validated. DiffSplice did not identify significant alternative splicing in any of these genes.

Calculation of the splice dissimilarity measure in SEVA makes this algorithm unique in being able to both quantify and visualize the relative heterogeneity of gene isoform usage in each phenotype. In Fig 4, we create multi-dimensional scaling (MDS) plots of the splice dissimilarity measure of the significant genes in SEVA. The closer two samples in the MDS plot, the less variable their junction expression profiles. As a result, these figures enable us to visually test the hypothesis that the modified Kendall-*τ* distance enables SEVA to identify more variable gene isoform usage in tumor than normal samples. The differentially spliced genes identified with SEVA (*DST, LAMA3, RASIP1*, and *TP63*) confirm that the variation of gene isoform expression in normal samples is lower than the variation in tumor samples, as hypothesized, and therefore not significant in EBSeq (Fig 4). On the other hand, *VEGFC* has consistent variability in cancer and normal samples and is not detected by SEVA (Supplemental Fig4). Therefore, SEVA is ideally suited to detect genes with more variable gene isoform usage in tumor relative to normal samples as hypothesized.

**Figure 4:**
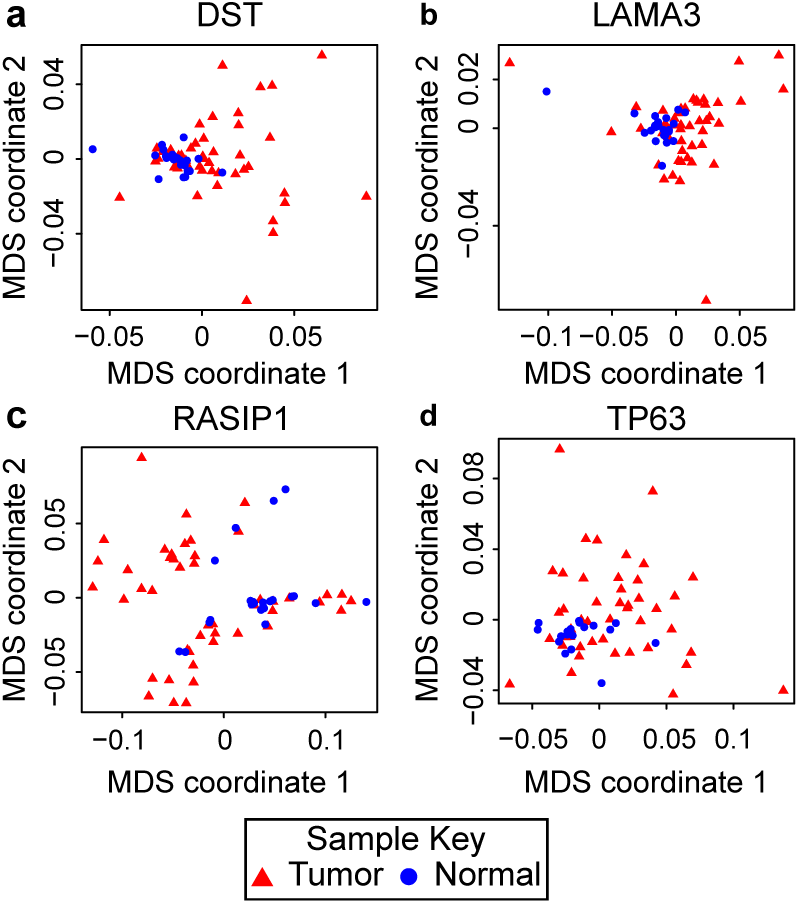
Multidimensional scaling (MDS) plot of splice dissimilarity measures in real HPV+ HNSCC junction expression from RNA-seq for (a) *DST*, (b) *LAMA3*, (c) *RASIP1*, and (d) *TP63*. Relative spread of samples in the MDS plots indicates their relative variability in normal samples (blue circles) and tumor samples (red triangles).

### 4.4 SEVA candidates in the training set are significantly enriched in cross-study validation on TCGA data

We also apply SEVA to independent RNA-seq data for 44 HPV+ HNSCC and 16 normal samples in TCGA (Cancer Genome Atlas Network, 2015) to cross-validate the ASE candidates in the training data from Guo *et al.*, 2016. 46% (352 out of 771 the gene candidates) of the hits are statistically significant in the SEVA analysis of the TCGA data. 214 of the genes identified on the training set are not expressed on the TCGA set. To test the significance of the list of genes and consistency of SEVA across two datasets, we check whether the ASE candidates from the training set are significantly enriched on the TCGA data. To do so, we calculate the SEVA p-values for all genes on the TCGA test set. A mean-rank geneset analysis indicates that the candidate genes identified on training are enriched among all genes with p-value < 2 × 10^−16^.

### 4.5 SEVA identifies more heterogeneity in splice variant usage in HPV-HNSCC samples with mutations in splice machinery genes

HNSCC tumors have two predominant subtypes: HPV+ and HPV-. HPV-tumors have greater genomic heterogeneity than HPV+ (Cancer Genome Atlas Network, 2015; Mroz *et al.*, 2015). Therefore, we apply SEVA to test whether there is higher inter-tumor heterogeneity in splice variant usage in these tumor subgroups. SEVA observes many genes as having alternative gene isoform usage between the two HNSCC subtypes,with little difference in relative heterogeneity (478 genes in HPV-and 338 in HPV+, Fig 5a). This finding occurs although a larger number of samples HPV-tumors (44 of 243, 18%) have alterations to genes in the splice variant machinery in contrast to HPV+ (3 of 36 with sequencing data, 8%). We further apply SEVA to compare inter-tumor heterogeneity of gene isoform usage in HPV-tumors with and without genetic alterations to the splicing machinery. We observe far fewer significant genes than in this comparison (Fig 5b). Nonetheless, the significant genes from this analysis have greater variability in samples with alterations in the splice variant machinery. These findings suggest that although the mechanisms of tumorigenesis are vastly different in HPV+ and HPV-tumors, both have similar heterogeneity in gene isoform usage. The mechanisms that cause mutations in the genes that encode for the splice variant machinery further increase the heterogeneity of splice variant usage in HPV-HNSCC.

**Figure 5:**
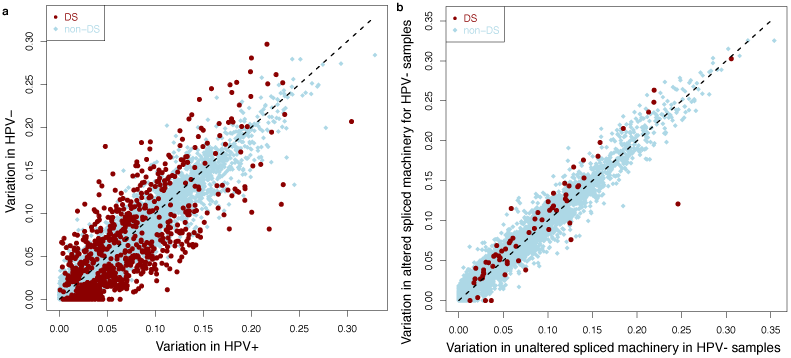
Comparison of differential gene isoform in TCGA HNSCC RNA-seq data. (a) Variability of junction expression profiles for genes significantly DS (red) from SEVA in HPV+vs HPV-HNSCC, respectively and not significantly DS (blue). (b) As for (a) comparing HPV-samples with and without alterations in RNA splice machinery genes.

## 5 Discussion

In this study, we develop SEVA to identify changes in the distribution of isoform expression in tumor samples to quantify heterogeneous gene isoform usage than that of normal samples. This SEVA algorithm has three benefits. First, focusing on the comparison of expression in overlapping junction pairs is computationally simpler than formulating a complete splicing graph that delineates all annotated gene isoforms (Song *et al.*, 2016; Hu *et al.*, 2013; Xing *et al.*, 2006). Second, the rank-based dissimilarity based upon Kendall-*τ* is blind to such coordinated changes in isoform expression that arise from differential expression. If the expression of a gene is higher in one sample than another, all its junctions will have higher expression, but their ranks will not change. Therefore, SEVA does not require normalization of isoform expression values based upon total gene expression that is used in other methods for differential splice variant analysis (Anders *et al.*, 2012; Guo *et al.*, 2017). Third, SEVA relies on junction expression and therefore can be applied to RNA-seq data processed with any alignment and quantification software.

We demonstrate the suitability of SEVA for differential heterogeneity of gene isoform usage by applying it to real and simulated RNA-seq data. Validation is performed against the modeled ASEs in simulated data. For real data, validation is performed with cross-study comparison of results from two cohorts of RNA-seq data for head and neck squamous cell carcinoma (HNSCC) tumors and normal samples. We also apply SEVA to benchmark its performance for experimentally validated splice variants of HNSCC from a previous microarray study (Li *et al.*, 2014). We further apply SEVA to perform a novel analysis of differential heterogeneity of gene isoform usage between the dominant HNSCC subtypes (HPV+ and HPV-) and between HPV-HNSCC samples with and without mutations to genes that regulate the splice variant machinery. Together, the results of these analyses in simulated and real data show that SEVA is a robust algorithm for inter-tumor heterogeneity in gene isoform usage in cancer samples relative to samples from a control group.

Consistent with the formulation of SEVA to detect differential heterogeneity of gene isoform usage between sample groups, we observe that SEVA has higher precision than EBSeq (Leng *et al.*, 2013), DiffSplice (Hu *et al.*, 2013), or rMATS (Shen *et al.*, 2014) in simulated datasets that model greater heterogeneity among tumor samples than among normal samples. The precision of SEVA, DiffSplice, and rMATS remain independent of the heterogeneity of gene isoform usage in the tumor samples, whereas that of EBSeq decreases with increasing homogeneity in gene isoform usage in the tumor samples. While SEVA retains a lower false positive rate in the simulated data, the recall depends on the heterogeneity of gene isoform usage. In our simulations, as the ratio of disrupted samples in the cancer batch increases, the recall of SEVA reduces dramatically (from 70% to 20%). DiffSplice and rMATS show almost constant recall (around 40-50% and 50-60%, respectively). While EBSeq recall increases with the homogeneity in gene isoform usage, SEVA loses its recall when gene isoform usage is greater than 80%. SEVA performs relatively best in the case of high heterogeneity of junction expression in the tumor population. Notably, as the number of cancers with an ASE increases the junction expression profiles are more homogeneous and therefore not accurately detected with SEVA. We hypothesize that SEVA will have lower recall than techniques based upon differential isoform expression in populations with homogeneous isoform usage. However, cancer samples are more heterogeneous and encompass a bigger spectrum of subtypes (Afsari *et al.*, 2014a; Corrada Bravo *et al.*, 2012; Eddy *et al.*, 2010). In practice, we hypothesize that differentially spliced genes show multiple patterns of isoform expression in tumors in multiple different cancer subtypes. Therefore, we anticipate far less than 80% homogeneity in driver splice events in cancer. Based upon the simulated data and pervasive genomic heterogeneity in tumors, we hypothesize that SEVA is uniquely suited to identify clinically relevant gene isoform usage in tumors and their subtypes. Moreover, it also the only algorithm that can quantify the extent of such heterogeneity of isoform usage for each gene.

The formulation of SEVA to detect differential heterogeneity of gene isoform usage between sample groups is unique. Notably, SEVA seeks genes with different distributions of gene isoform usage among samples within the same and between distinct phenotypes than distributions sought by algorithms for differential gene isoform expression, including EBSeq (Leng *et al.*, 2013), DiffSplice (Hu *et al.*, 2013), and rMATS (Shen *et al.*, 2014). Care must be taken in interpreting the comparisons between methods that are presented. These comparisons do not demonstrate that one method outperforms another in accuracy, but rather to benchmark the dependency of their performance on gene isoform usage. The simulation studies highlight that SEVA requires differential heterogeneity of gene isoform usage to infer candidates, as anticipated in the algorithm’s design. Similarly, EBSeq is designed to detect differential isoform abundance and not differential splicing. Therefore, if a gene’s abundance changes across conditions, even if the relative (within-gene) frequencies of the isoforms of a gene remain the same (no differential splicing), EBSeq would consider all of the isoforms to have differential abundance. Thus, different methods will find distinct alternative splicing events. Characterizing the events modeled with all these techniques is essential to characterize transcriptional diversity and the distinct contributions of alternative splicing to biological function.

SEVA, EBSeq, and DiffSplice could all be applied to RNA-seq data normalized with MapSplice (Wang *et al.*, 2010) to enable cross-study validation of the data from Guo *et al.* 2016 with the TCGA normalized data. However, rMATS could not be applied to MapSplice aligned data. While there are numerous other algorithms for such differential splice analysis, many rely on data obtained from distinct alignment and normalization pipelines (Anders *et al.*, 2012; Shen *et al.*, 2012; Song *et al.*, 2016; Guo *et al.*, 2017). These preprocessing techniques may introduce additional variables into the differential splice variant analysis, complicating the direct comparisons of gene candidates on *in silico* and RNA-seq datasets presented in this paper. Notably, SEVA inputs junction expression to use direct evidence of alternative splice usage, including intron retention. Therefore, the algorithm can be readily applied to junction expression obtained from other aligners. It is also directly applicable to estimates of percent spliced (Alamancos *et al.*, 2015) or transcript-level expression in place of junction expression, which can be compared in future studies. Therefore, future studies are needed to compare the performance of such differential splice variant algorithms across normalization pipelines on real biological datasets with known ground truth of gene isoform usage. Nonetheless, the SEVA algorithm is applicable for differential splice variant analysis from junction expression from any alignment algorithm and its rank-based statistics make it likely to be independent of the normalization procedure (Afsari *et al.*, 2014b; Bolstad *et al.*, 2003).

The p-values for the rank-based SEVA statistics are computed using the normal approximation to U-statistics described previously (Afsari *et al.*, 2014a). This approximation yields computational efficiency, with the analysis of the 279 HNSCC tumor samples in the TCGA freeze set (Cancer Genome Atlas Network, 2015) completing in under 2 hours on a standard laptop processor. Thus, SEVA is readily applicable to splice variant analysis of large cohort studies. However, this formulation does introduce two weaknesses that must be addressed in future work. First, genes that have multiple overlapping junctions with zero read coverage will have ties that will introduce inaccuracies to the rank comparisons used in SEVA (Fig 1). We note that such ties are a common problem to all non-parametric algorithm. Such zero values are likely to be pervasive in lowly expressed genes because reads will have a low probability of covering all the junctions of a gene in each splice variant. We note that analysis of lowly expressed genes is challenging generally for RNA-seq analyses. To avoid this challenge, we constrained the simulated data in this study to genes with moderate to high expression. We relaxed this constraint in real data, and nonetheless observed significant cross-study concordance of inferred splice variants. We note that the software implementing SEVA contains several filtering parameters which may be selected by users to optimize performance on individual datasets. Future work is needed to handle the ties that result from such lowly expressed genes and to evaluate the filtering criterion to optimize the performance of SEVA in these cases. A second challenge to SEVA arises in the large sample-sizes required for analysis. Theoretical work shows that the normal approximation to U-statistics that are used in SEVA require a minimum of thirty samples (total in both phenotypes) for accurate estimation ofp-values (Vaart, 1998). Reducing the number of samples will yield an incorrect estimate of the covariance matrix used in SEVA, thereby underestimating the p-values obtained from the algorithm. Future work adapting empirical Bayes estimates of the covariance will be essential to extend SEVA to analyze heterogeneity of splice variant usage in smaller sample cohorts.

In simulations with 25 samples per group, SEVA has uniformly high precision relative to EBSeq and DiffSplice in detecting ASEs. Our simulations suggest that SEVA performs better in scenarios in which cancer samples have a higher degree of heterogeneity compared to normal samples consistent with the unique formulation of the method. As further validation, genes with alternative splicing events in HPV+ HNSCC from Guo *et al.*, 2016 were significantly enriched in cross-study validation on RNA-seqdata for HPV+ HNSCC samples in TCGA (Cancer Genome Atlas Network, 2015). Moreover, the modified Kendall-*τ* dissimilarity metric used in SEVA also accurately characterizes the higher heterogeneity of gene isoform usage in tumors relative to normal in the confirmed HNSCC-specificASEs DST, LAMA3, TP63, and RASIP1 identified in previous microarray analysis (Li *et al.*, 2014). As further justification for our simulation, we observed that the MDS visualization of the heterogeneity of gene isoform for these genes resembles the same MDS visualization of alternatively spliced genes in the *in silico* data. These data show that SEVA is adept at inferring ASEs in tumor samples with heterogeneous gene isoform usage relative to normal samples.

SEVA is uniquely designed to quantify which phenotype has more variable gene isoform usage. The algorithm observes higher inter-tumor heterogeneity in splice variant usage in HPV+ HNSCC tumors relative to normal samples, consistent with the hypothesis of inter-tumor heterogeneity that led to the development of the algorithm. HNSCC is divided into two primary subtypes (HPV-and HPV+). Of these, HPV-tumors are established as having more variable genetic alterations than HPV+ tumors (Cancer Genome Atlas Network, 2015; Mroz *et al.*, 2015). Nonetheless, SEVA analysis identifies little difference in the heterogeneity of splice variant usage between the tumor types (478 genes with differential heterogeneity of alternative splicing in HPV-and 338 in HPV+). This similarity is observed despite different samples sizes (44 HPV+ and 235 HPV-), suggesting that the SEVA statistics are robust to imbalanced study design. In addition, some HPV-samples have genetic alterations in genes that regulate the splice variant machinery. SEVA identifies that HPV-HNSCC tumors with alterations in these genes have greater variation in isoform usage than those that do not (52 genes with differential heterogeneity of alternative splicing in samples with broken machinery and 14 genes in samples without). Together, these analyses suggest that SEVA can distinguish differences in heterogeneity in isoform usage that are specific to cancer subtypes. Moreover, SEVA observes greater variation in splice variant usage in tumor samples with genetic alterations to the splice variant machinery within a specific cancer subtype. Further pan-cancer and pan-genomics are essential to quantify the heterogeneity of gene isoform usage across cancers and their subtypes.

## Acknowledgements

We thank Simina Boca, Leslie Cope, Ludmila V Danilova, Sarah Wheelan, and members of New PI Slack for advice on analyses, data preprocessing, data access, and suggestions.

